# Accurate prediction of single-cell DNA methylation states using deep learning

**DOI:** 10.1101/055715

**Authors:** Christof Angermueller, Heather J. Lee, Wolf Reik, Oliver Stegle

## Abstract

Recent technological advances have enabled assaying DNA methylation at single-cell resolution. Current protocols are limited by incomplete CpG coverage and hence methods to predict missing methylation states are critical to enable genome-wide analyses. Here, we report DeepCpG, a computational approach based on deep neural networks to predict DNA methylation states from DNA sequence and incomplete methylation profiles in single cells. We evaluated DeepCpG on single-cell methylation data from five cell types generated using alternative sequencing protocols, finding that DeepCpG yields substantially more accurate predictions than previous methods. Additionally, we show that the parameters of our model can be interpreted, thereby providing insights into the effect of sequence composition on methylation variability.

## Background

DNA methylation is one of the most extensively studied epigenetic marks, and is known to be implicated in a wide range of biological processes, including chromosome instability, X-chromosome inactivation, cell differentiation, cancer progression and gene regulation [1–4].

Well-established protocols exist for quantifying average DNA methylation levels in populations of cells. Recent technological advances have enabled profiling DNA methylation at single-cell resolution, either using genome-wide bisulfite sequencing (scBS-seq [5]) or reduced representation protocols (scRRBS-seq [6–8]). These protocols have already provided unprecedented insights into the regulation and the dynamics of DNA methylation in single cells [6, 9], and have uncovered new linkages between epigenetic and transcriptional heterogeneity [8, 10, 11].

Because of the small amounts of genomic DNA starting material per cell, single-cell methylation analyses are intrinsically limited by moderate CpG coverage (Figure 1a, 20-40% for scBS-seq [5]; 1-10% for scRRBS-seq [6–8]). Consequently, a first critical step is to predict missing methylation states to enable genome-wide analyses. While methods exist for predicting average DNA methylation profiles in cell populations [12–16], these approaches do not account for cell-to-cell variability. Additionally, existing methods require *a priori* defined features and genome annotations, which are typically limited to a narrow set of cell types and conditions.

**Figure 1.**
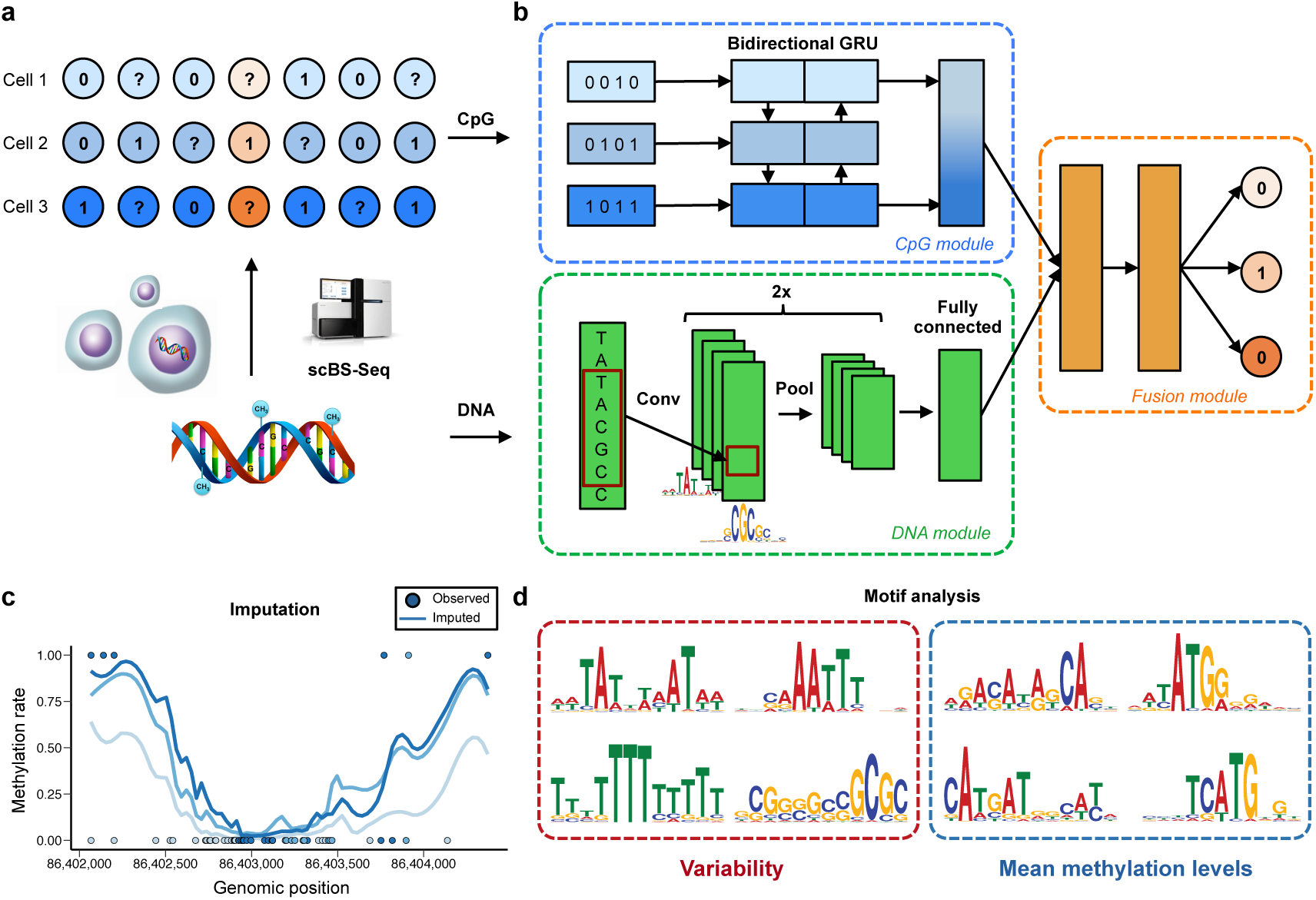
DeepCpG model training and applications. (a) Sparse single-cell CpG profiles, for example as obtained from scBS-seq [5] or scRRBS-seq [6–8]. Methylated CpG sites are denoted by ones, un-methylated CpG sites by zeros, and question marks denote CpG sites with unknown methylation state (missing data). **(b)** Modular architecture of DeepCpG. The DNA module consists of two convolutional and pooling layers to identify predictive motifs from the local sequence context, and one fully connected layer to model motif interactions. The CpG module scans the CpG neighbourhood of multiple cells (rows in **b**), using a bidirectional gated recurrent network (GRU [24]), yielding compressed features in a vector of constant size. The fusion module learns interactions between higher-level features derived from the DNA- and CpG module to predict methylation states in all cells. **(c, d)** The trained DeepCpG model can be used for different downstream analyses, including genome-wide imputation of missing CpG sites **(c)** and the discovery of DNA sequence motifs that are associated with DNA methylation levels or cell-to-cell variability **(d)**.

Here, we report DeepCpG, a computational method based on deep neural networks [17–19] for predicting single-cell methylation states and for modelling the sources of DNA methylation variability. DeepCpG leverages associations between DNA sequence patterns and methylation states as well as between neighbouring CpG sites, both within individual cells and across cells. Unlike previous methods [12, 13, 15, 20–23], our approach does not separate the extraction of informative features and model training. Instead, DeepCpG is based on a modular architecture and learns predictive DNA sequence- and methylation patterns in a data-driven manner. We evaluated DeepCpG on mouse embryonic stem cells profiled using whole-genome single-cell methylation profiling (scBS-seq [5]), as well as on human and mouse cells profiled using a reduced representation protocol (scRRBS-seq [8]). On all cell types, DeepCpG yielded substantially more accurate predictions of methylation states than previous approaches. Additionally, DeepCpG uncovered both previously known and de novo sequence motifs that are associated with methylation changes and methylation variability between cells.

## Results and discussion

DeepCpG is trained to predict binary CpG methylation states from local DNA sequence windows and observed neighbouring methylation states (Figure 1a). A major feature of the model is its modular architecture, consisting of a CpG module to account for correlations between CpG sites within and across cells, a DNA module to detect informative sequence patterns, and a fusion module that integrates the evidence from the CpG and DNA module to predict methylation states at target CpG sites (Figure 1b).

Briefly, the DNA and CpG module were designed to specifically model each of these data modalities. The DNA module is based on a convolutional architecture, which has been successfully applied in different domains [25–28], including genomics [29–33]. The module takes DNA sequences in windows centred on target CpG sites as input, which are scanned forsequence motifs using convolutional filters, analogous to conventional position weight matrices [34, 35] (**Methods**). The CpG module is based on a bidirectional gated recurrent network [24], a sequential model that compresses patterns of neighbouring CpG states from a variable number of cells into a fixed-length feature vector (**Methods**). Finally, the fusion module learns interactions between output features of the DNA- and CpG module, and predicts the methylation state at target sites in all cells using a multi-task architecture. The trained DeepCpG model can be used for different downstream analyses, including i) to impute low-coverage methylation profiles for sets of cells (Figure 1c), and ii) to discover DNA sequence motifs that are associated with methylation states and cell-to-cell variability (Figure 1d).

### Accurate prediction of single-cell methylation states

First, we assessed the ability of DeepCpG to predict single-cell methylation states and compared the model to existing imputation strategies for DNA methylation (**Methods**). As a baseline approach, we considered local averaging of the observed methylation states, either in 3 kb windows centred on the target site of the same cell (WinAvg) [36], or across cells at the target site (CpGAvg). Additionally, we compared DeepCpG to a random forest classifiers [37] trained on individual cells using the DNA sequence information and neighbouring CpG states as input (RF). Finally, we evaluated a recently proposed random forest model to predict methylation rates for bulk ensembles of cells [12], which takes comprehensive DNA annotations into account, including genomic contexts, and tissue-specific regularly annotations such as DNase1 hypersensitivity sites, histone modification marks, and transcriptionfactor binding sites (RF Zhang). All methods were trained, selected, and tested on distinct chromosomes via holdout validation (**Methods**). Since the proportion of methylated versus unmethylated CpG sites can be unbalanced in globally hypo- or hyper-methylated cells, we used the area under the receiver operating characteristics curve (AUC) to quantify the prediction performance of different models. We have also considered a range of alternative metrics, including precision-recall curves, F1 score [38], and Matthews correlation coefficient [39], resulting in overall consistently conclusions (**Additional File 1:**Figure 1–Figure 3).

Initially, we applied all methods to 18 serum-cultured mouse embryonic stem cells (mESCs, average CpG coverage 17.7%, **Additional File 1:** Figure 4), profiled using whole-genome single-cell bisulfite sequencing (scBS-seq [5]).

DeepCpG yielded more accurate predictions than any of the alternative methods, both genome-wide and in different genomic contexts (Figure 2). Notably, DeepCpG was consistently more accurate than RF Zhang, a model that relies on genomic annotations. These results indicate that DeepCpG can automatically learn higher-level annotations from the DNA sequence. This ability is particularly important for analysing single-cell datasets, where individual cells may be from different cell types and states, making it difficult to derive appropriate annotations.

**Figure 2.**
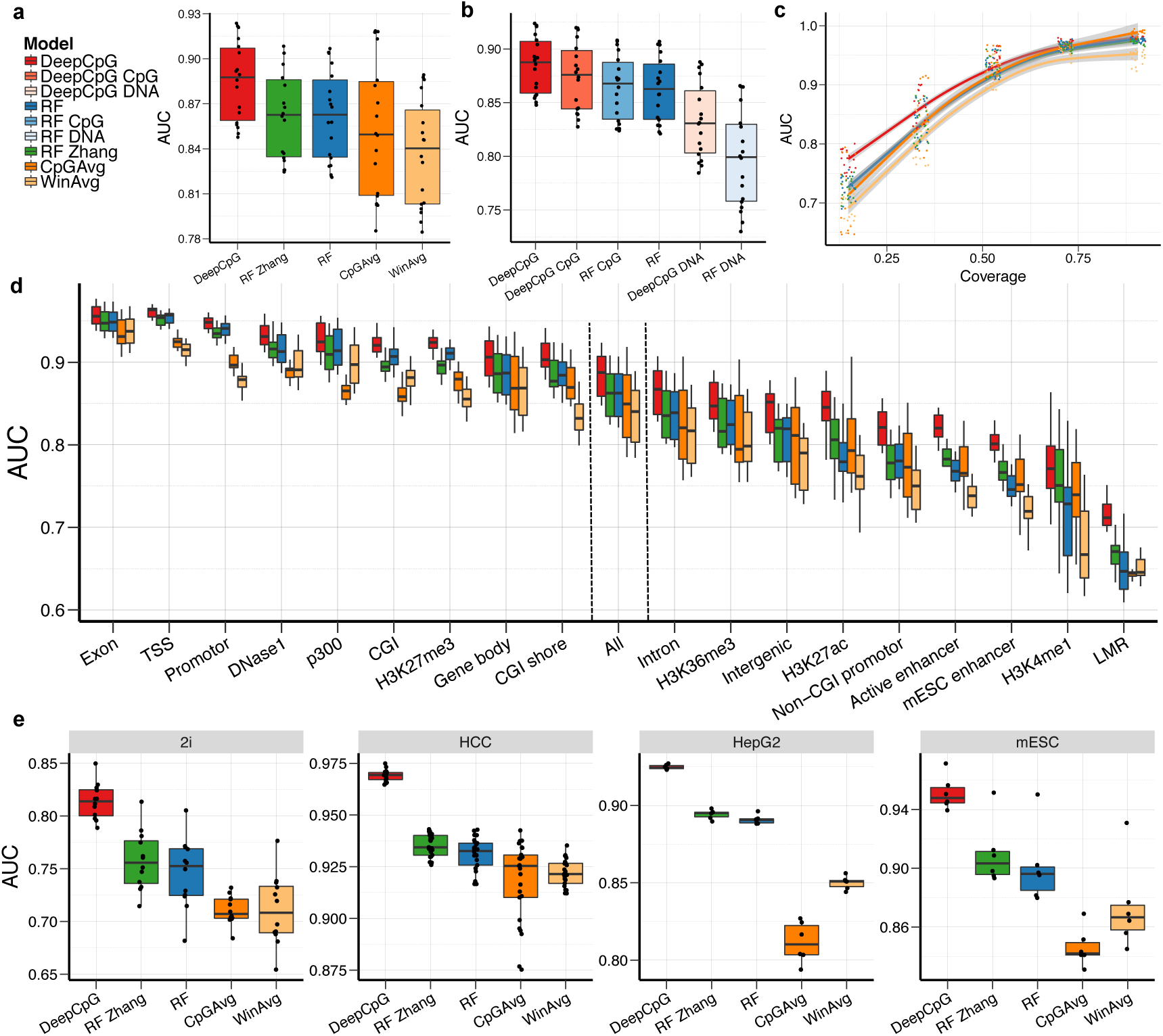
DeepCpG accurately predicts single-cell CpG methylation states. (a) Genome-wide prediction performance for imputing CpG sites in 18 serum-grown mouse embryonic stem cells (mESCs) profiled using scBS-seq [5]. Performance is measured by the area under the receiver-operating characteristic curve (AUC), using holdout validation. Considered were DeepCpG, a random forest classifiers trained either using DNA sequence and CpG features (RF), or trained using additional annotations from corresponding cell types (RF Zhang [12]). Baseline models inferred missing methylation states by averaging observed methylation states, either across consecutive 3 kb regions within individual cells (WinAvg [5]), or across cells at a single CpG site (CpGAvg). (**b**) Performance breakdown of DeepCpG and RF, considering models either trained exclusively using methylation features (DeepCpG CpG, RF CpG) or DNA features (DeepCpG DNA, RF DNA). (**c**) AUC of the models as in (**a**) stratified by genomic contexts with variable coverage across cells. Trend lines were fit to the observed coverage levels using local polynomial regression (LOESS [40]); shaded areas denote 95% confidence intervals. (**d**) AUC for alternative sequence contexts. *All* corresponds to the genome-wide performances as in (**a**). (**e**) Genome-wide prediction performance on 12 2i-grown mESCs profiled using scBS-seq [5], as well as three cell types profiled using scRRBS-seq [8], including 25 human HCC cells, 6 HepG2 cells, and 6 additional mESCs.

To assess the relative importance of DNA sequence features compared to neighbouring CpG sites, we trained the same models, however, either exclusively using DNA sequence features (DeepCpG Seq, RF Seq) or neighbouring methylation states (DeepCpG CpG, RF CpG). Consistently with previous studies in bulk populations [12], methylation states were more predictive than DNA features, and models trained with both CpG and DNA features performed best (Figure 2b). Notably, DeepCpG trained with CpG features alone outperformed a random forest classifiers trained with both CpG and DNA features. A likely explanation for the accuracy of the CpG module is its recurrent network architecture, which enables the module to effectively transfer information from neighbouring CpG sites across different cells (**Additional File 1: Figure 5, 20**).

The largest relative gains between RF and DeepCpG were observed when training both models with DNA sequence information only (AUC 0.83 versus 0.80, Fig. 2b). This demonstrates the strength of the DeepCpG DNA module to extract predictive sequence features from wide DNA sequence windows of up to 1001 bp (**Additional File 1: Figure 7a**), which is in particular critical for accurate predictions from DNA in uncovered genomic regions, for example when using reduced representation sequencing data [6–8]. Consistent with this, DeepCpG outperformed other methods by a large margin in genomic contexts with low CpG coverage (Figure 2c, **Additional File 1: Figure 6**).

Next we explored the prediction performance of all models in different genomic contexts. In line with previous findings [12, 13], all models performed best in GC-rich contexts (Figure 2d). However, the relative gains in performance of DeepCpG were largest in GC-poor genomic contexts, including non-CGI promoters, enhancer regions, and histone modification marks (H3K4me1, H3K27ac) — contexts that are known to be associated with higher methylation variability between cells [36].

We also applied DeepCpG to 12 2i-cultured mESCs profiled using scBS-seq [5] and to data from three cell types profiled using scRRBS-seq [8], including 25 human hepatocellular carcinoma cells (HCC), 6 human heptoplastoma-derived (HepG2) cells, and an additional set of 6 mESCs. Notably, in contrast to the serum cells, the human cell types are globally hypo-methylated (**Additional File 1:** Figure 4). Across all cell types, DeepCpG yielded substantially more accurate predictions than alternative methods, demonstrating the broad applicability of the model, including to hypo- and hyper-methylated cells, as well to data generated using different sequencing protocols.

### Estimation of the effect of DNA motifs and sequence mutations on methylation states

In addition to imputing missing methylation states, DeepCpG can be used to discover methylation-associated motifs, and to investigate the effect of DNA sequence mutations on CpG methylation.

To explore this, we used the DeepCpG DNA module trained on serum mESCs, and analysed the learnt filters of the first convolutional layer. These filters recognize DNA sequence motifs similarly to conventional position weight matrices, and can be visualised as sequence logos (Figure 3, **Additional File 2**). We considered two complementary metrics to assess the importance of the 128 motifs discovered by DeepCpG: i) their occurrence frequency in DNA sequence windows (activity), and ii) their estimated association with single-cell methylation states (**Additional File 1: Figure 8**). To investigate the co-occurrence of motifs across sequence windows, we applied principal component analysis (Figure 3) and hierarchical clustering (**Additional File 1: Figure 9**, **10**) to motif activities.

**Figure 3.**
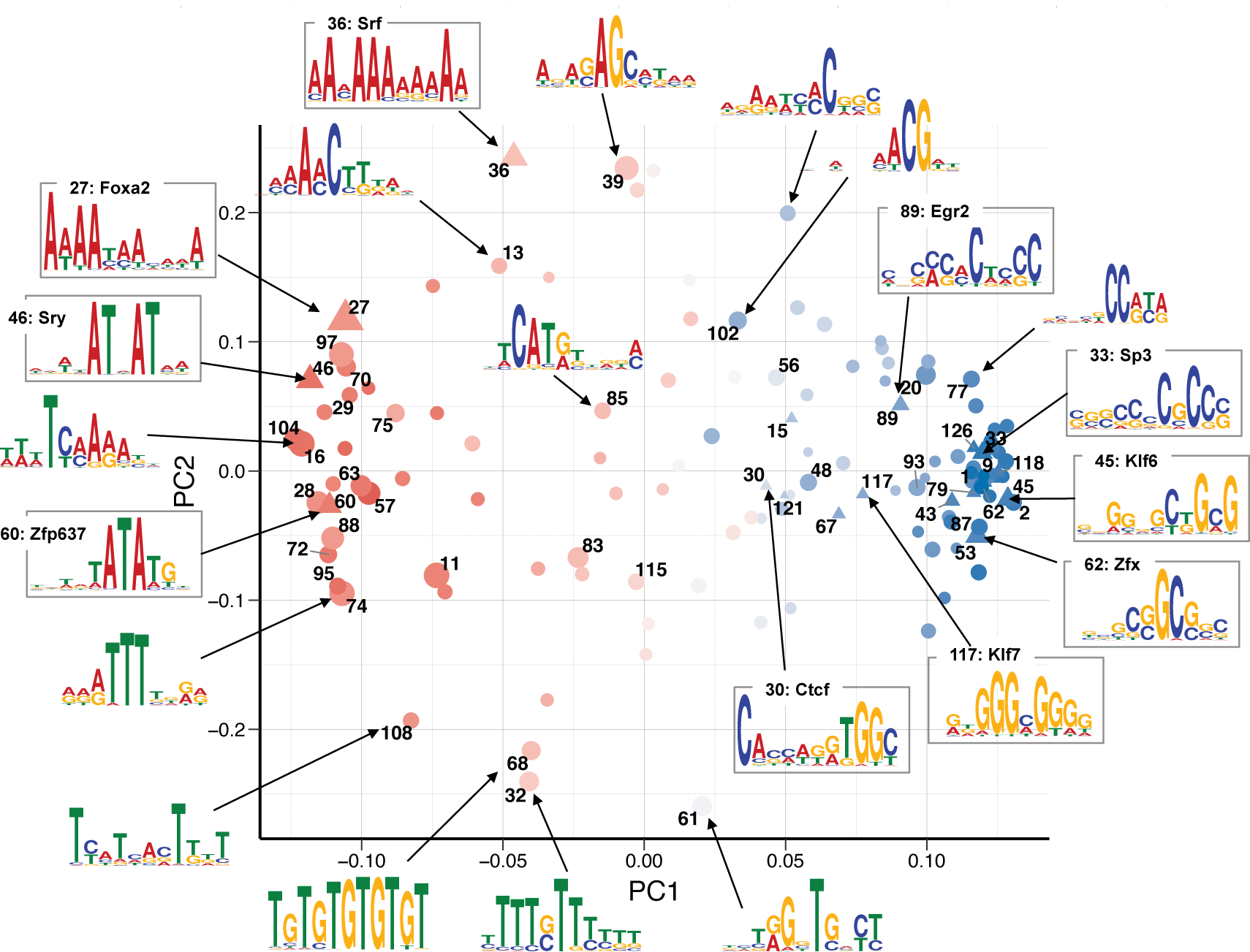
Discovered sequence motifs associated with DNA methylation. Clustering of 128 motifs discovered by DeepCpG. Shown are the first two principal components of the motif occurrence frequencies in sequence windows (activity). Triangles denote motifs with significant (FDR<0.05) similarity to annotated motif in the CIS-BP [43] or UniPROPE [44] database. Marker size indicates the average activity; the estimated motif effect on methylation level is shown in colour. Sequence logos are shown for representative motifs with larger effects, including 10 annotated motifs

Motifs with similar nucleotide composition tended to co-occur in the same sequence windows, where two major motif clusters were associated with increased or decreased methylation levels (**Additional File 1: Figure 11**). Consistent with previous findings [16, 41, 42], we observed that motifs associated with decreased methylation tended to be CG rich and were most active in CG rich promoter regions, transcription start sites, as well as in contexts with active promoter marks such as H3K4me3 and p300 sites (**Additional File 1: Figure 10**). Conversely, motifs associated with increased methylation levels tended to be AT rich and were most active in CG poor genomic contexts (**Additional File 1: Figure 10**).

20 out of the 128 discovered motifs significantly (FDR<0.05) matched motifs annotated in the CIS-BP [43] and UniPROPE [44] database. 17 of these motifs were transcription factors with a known implication in DNA methylation [16, 45, 46], including CTCF [47], E2f [48], and members of the Sp/KLF family [49]— transcription factors and regulators of cell differentiation. 13 out of the 20 annotated motifs had been shown to interact with DNMT3a and DNMT3b [45], two major DNA methylation enzymes. Three motifs have no clear associations with DNA methylation. These included Foxa2 [50, 51] and Srf [52, 53], which play roles in cell differentiation and embryonic development, as well as Zfp637 [54, 55], a zinc finger protein that had recently been implicated with spermatogenesis in mouse.

The trained DeepCpG model can also be used to estimate the effect of single nucleotide mutations on CpG methylation. In order to efficiently assess the mutational effect, we adapted a gradient-based approach [56], which is markedly more efficient than previous approaches [30, 31, 33] (**Methods**). As expected, sequence changes in the direct vicinity of the target site had the largest effects (Figure 4). Mutations in CG dense regions such as CpG islands or promoters tended to have smaller effects, suggesting that DNA methylation in these genomic contexts is more robust to single base-pair mutations. Globally, we observed a negative correlation between the predicted effect of single-nucleotide changes and DNA sequence conservation (P < 1.0×10^−15^, **Additional File 1: Figure 12**), providing evidence that estimated mutational effects capture genuine effects. We further investigated DeepCpG effect predictions in HepG2 cells for 2,379 methylation QTLs (mQTLs) [57], and found that effects are significantly larger for known mQTLs compared to matched random variants (P < 1.0×10^-15^, Wilcoxon rank sum test; **Additional File 1: Figure 13, 14**).

**Figure 4.**
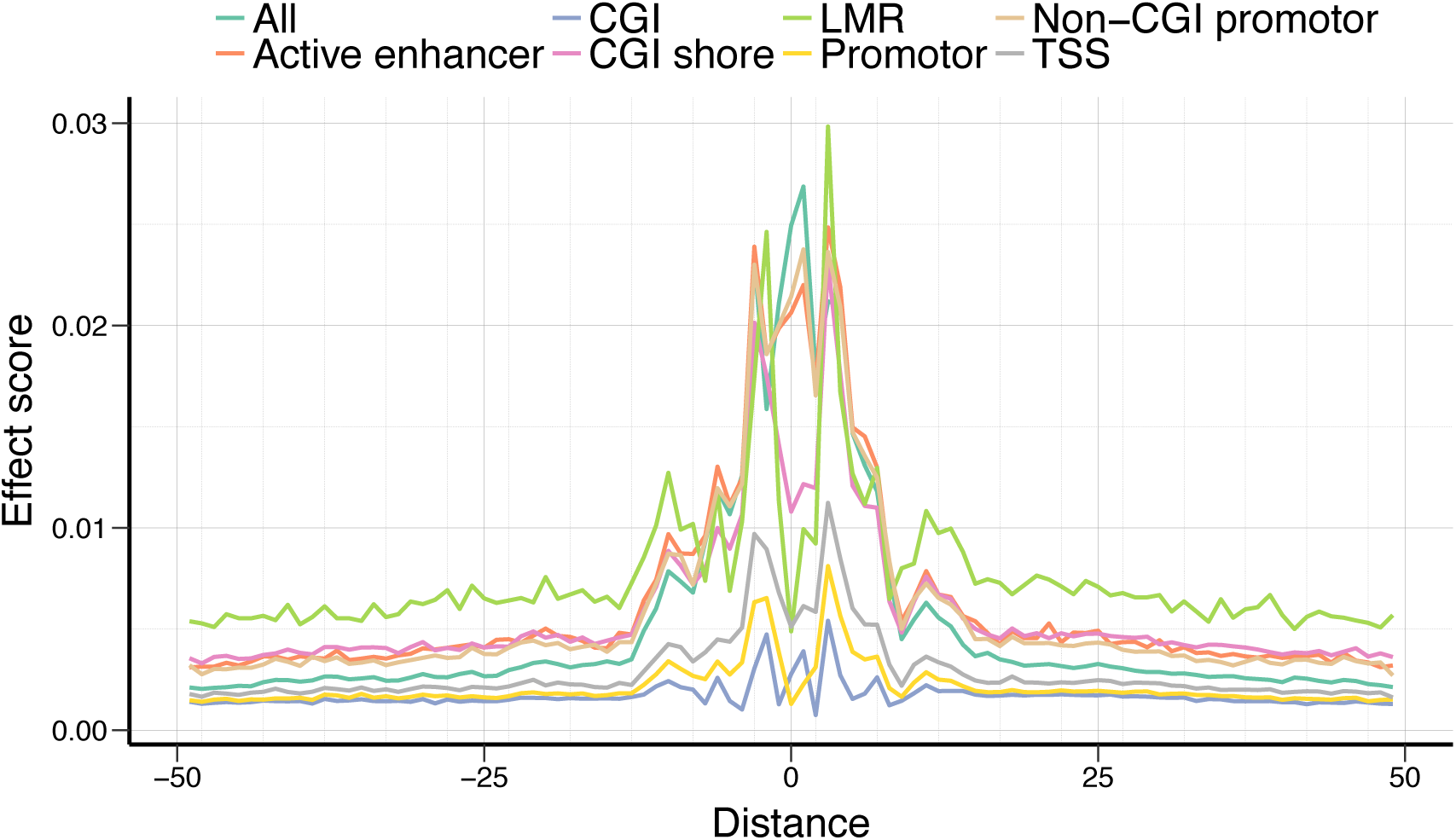
Effect of point mutations on DNA methylation. Average genome-wide predicted effect of DNA sequence mutations in different genomic contexts as a function of the distance to the CpG site.

### Discovery of DNA motifs that are associated with methylation variability

We further analysed the influence of motifs discovered by DeepCpG on methylation variability between cells.

To discern motifs that affect variability between cells from those that affect the average methylation level, we trained a second neural network. This network had the same architecture and in particular reused the motifs from the DNA module of DeepCpG, however was trained to jointly predict the variability across cells and the mean methylation level of each CpG site (**Methods**).

Notably, this model could predict both global changes in mean methylation levels (Pearson’s R=0.80, MAD=0.01, **Additional File 1: Figure 15**), as well as cell-to-cell variability (Pearson’s R=0.44, MAD=0.03, Figure 5d; Kendall’s R=0.29, **Additional File 1: Figure 16**).

**Figure 5.**
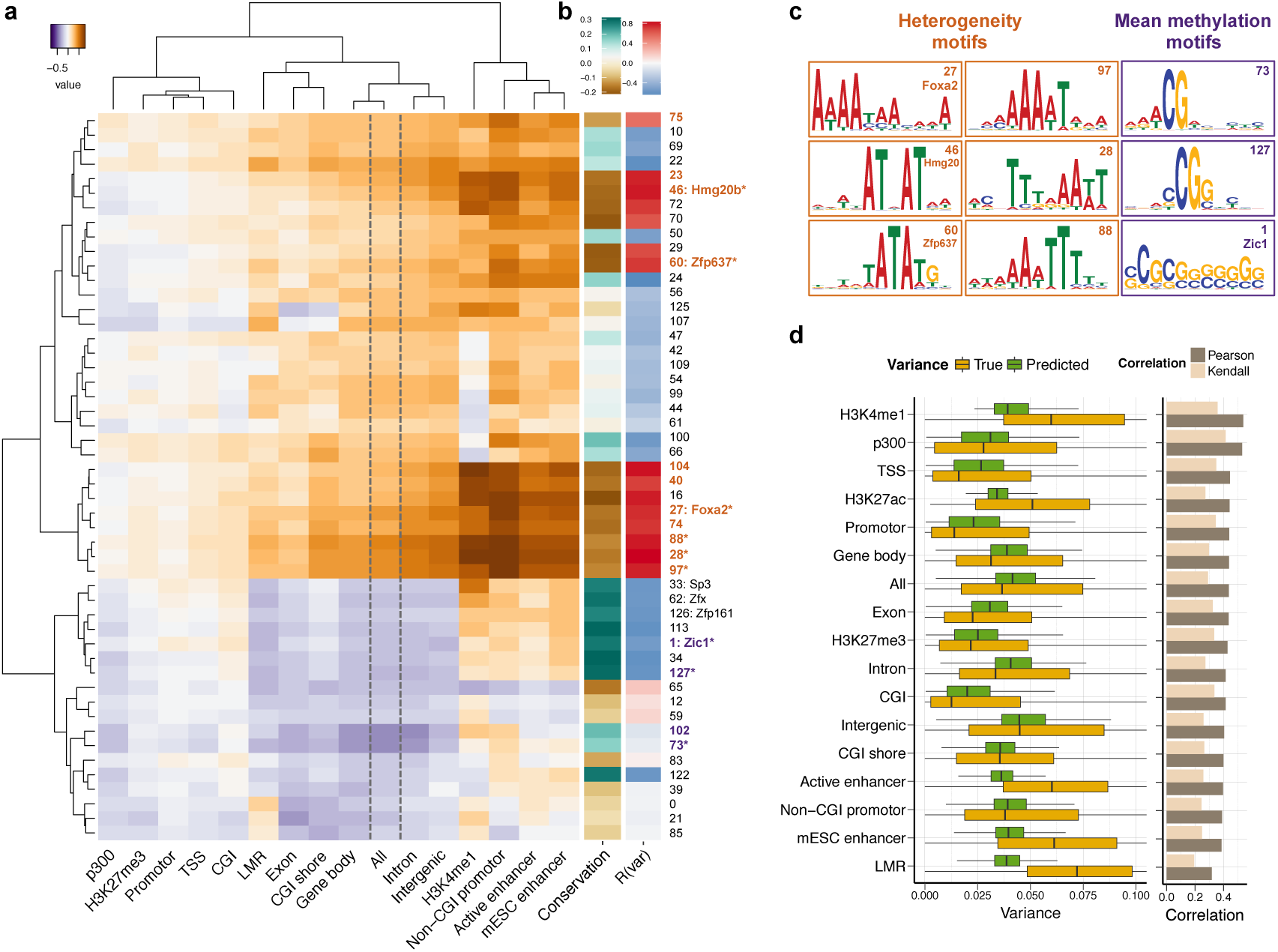
Prediction of methylation variability from local DNA sequence. (a) Difference of motif effect on cell-to-cell variability and methylation levels for different genomic contexts on test chromosomes. Motifs associated with increased cell-to-cell variability are highlighted in brown; motifs that were primarily associated with changes in methylation level are shown in purple. (**b**) Genome-wide correlation coefficients between motif activity and DNA sequence conservation (left), as well as cell-to-cell variability (right). (**c**) Sequence logos for selected motifs identified in (**a**), which are marked by asterisks in (**b**). (**d**) Boxplots of the predicted and the observed cell-to-cell variability for different genomic contexts on held out test chromosomes (left), alongside Pearson’s and Kendall’s correlation coefficients within contexts (right).

In general, there is an intrinsic mean-variance relationship of single-cell methylation states (**Additional File 1: Figure 17**), and hence the separation of the motif impact on mean methylation and methylation variance is partially confounded. To disentangle this relationship, we developed an approach to separately estimate the effect of individual motifs on cell-to-cell variability and mean methylation levels (**Methods**). Briefly, we quantified motif effects by correlating motif activities with predicted mean methylation levels and cell-to-cell variability, and used the difference between these effect size estimates to identify variance- and mean methylation associated motifs. This analysis identified 22 motifs that were primarily associated with cell-to-cell variance (Figure 5). These motifs were most active in CG-poor and active enhancer regions — sequence contexts with increased epigenetic variability between cells [36]. 12 of the identified motifs were AT-rich and associated with increased variability, including the differentiation factors Foxa2 [50, 51], Hmg20b [58], and Zfp637 [54, 55]. Notably, variance-increasing motifs were more frequent in un-conserved regions such as active enhancers, in contrast to variance- decreasing motifs, which were enriched in evolutionary conserved regions such as gene promoters (Figure 5b, **Additional File 1:Figure 18**). Our analysis also revealed 4 motifs that were primarily associated with mean methylation levels, which were in contrast CG rich and most active in conserved regions.

To explore whether the model predictions for variable sites are functionally relevant, we overlaid predictions with methylome-transcriptome linkages obtained using parallel single-cell methylation and transcriptome sequencing in the same cell type [10]. The rationale behind this approach is that regions with increased methylation variability are more likely to harbour associations with gene expression. Consistent with this hypothesis, we identified a weak but globally significant association (Pearson’s correlation R=0.11, *P*=5.72×10^−16^, **Additional File 1: Figure 19**).

## Conclusions

Here we reported DeepCpG, a computational approach based on convolutional neural networks for modelling low-coverage single-cell methylation data. In applications to mouse and human cells, we have shown that DeepCpG accurately predicts missing methylation states, and detects sequence motifs that are associated with changes in methylation levels and cell-to-cell variability.

We have demonstrated that our model enables accurate imputation of missing methylation states, thereby facilitating genome-wide downstream analyses. DeepCpG offers major advantages in shallow sequenced cells as well as in sparsely covered sequence contexts with increased methylation variability between cells. More accurate imputation methods may also help to reduce the required sequencing depth in single-cell bisulfite sequencing studies, thereby enabling the analysis of larger numbers of cells at reduced cost.

We have further shown that DeepCpG can be used to identify annotated and de novo sequence motifs that are predictive for DNA methylation levels or methylation variability, and to estimate the effect of DNA sequence mutations. Models such as DeepCpG further allow discerning pure epigenetic effects from variation that reflect DNA sequence changes. Although we have not considered this in our work, it would also be possible to use the model residuals for studying methylation variability that is unlinked to DNA sequence effects.

Finally, we have used additional data obtained from parallel methylation-transcriptome sequencing protocols [10] to annotate regions with increased methylation variability. An important area of future work will be to integrate multiple data modalities profiled in the same cells using parallel-profiling methods [8, 10], which are now becoming increasingly available for different molecular layers.

## Methods

**Accession codes** The scBS-seq data from 18 serum and 12 2i ES-cells have previously been described in Smallwood et al. *[5]* and are available under the Gene Expression Omnibus (GEO) accession number <u>GSE56879</u>.

**Availability of code** An implementation of DeepCpG is available at https://github.com/cangermueller/deepcpg.

## Acknowledgements

We are grateful to Yarin Gal for valuable discussions about quantifying prediction uncertainty using dropout. We are grateful to Leopold Parts for commenting on the manuscript, and Felix Krueger for pre-processing the data. O. S. is supported by the European Molecular Biology Laboratory (EMBL), the Wellcome Trust and the European Union.

## Author contributions

C.A. and O.S. devised and developed the model. C.A. implemented and evaluated the model. C.A., H.J.L., W.R. and O.S. interpreted the results. C.A. and O.S. wrote the paper.

## Author information

The authors declare no competing financial interests. Correspondence and requests for materials should be addressed to

## Abbreviations

AUC: Area under the receiver operating characteristic curve; bp: base pair; CGI: CpG island; CNN: Convolutional Neural Network; DHS: DNAse I hypersensitive; RF: random forest; ESC: embryonic stem cell; ROC: receiver operating characteristic; scBS-seq: single-cell bisulfite sequencing; scM&T: single-cell methylome and transcriptome sequencing; scRNA-seq: single-cell RNA sequencing; scRRBS-seq: single-cell reduced representation bisulfite sequencing; TFBS: transcription factor binding site; WinAvg: window averaging; WGBS: whole-genome bisulfite sequencing.

## DeepCpG model

DeepCpG consists of a DNA module to extract features from the DNA sequence, a CpG module to extract features from the CpG neighbourhood of all cells, and a multi-task fusion model that integrates the evidence from both modules to predict the methylation state of target CpG sites for multiple cells.

### DNA module

The DNA module is a convolutional neural network (CNN) with multiple convolutional and pooling layers, and one fully connected hidden layer. CNNs are designed to extract features from high-dimensional inputs while keeping the number of model parameters tractable by applying a series of convolutional and pooling operations. Unless stated otherwise, the DNA module takes as input a 1001 bp long DNA sequence centred on a target CpG site *n*, which is represented as a binary matrix *s_n_* by one-hot encoding the *D = 4* nucleotides as binary vectors A=[1, 0, 0, 0], T=[0,1, 0, 0], G=[0, 0,1, 0], and C=[0, 0, 0, 1]. The input matrix *s_n_* is first transformed by a 1d-convolutional layer, which computes the *activations a_nfi_* of multiple convolutional filters *f* at every position *i* as follows:

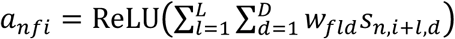

Here, *w_f_* are the parameters or *weights* of convolutional filter *f* length *L*. These can be interpreted similarly to position weight matrices (PWM), which are matched against the input sequence *s_n_* at every position *i* to recognize distinct motifs. The ReLU(*x*)= max(0,*x*) activation function sets negative values to zero, such that *a_nfi_* corresponds to the evidence that the motif represented by *w_f_* occurs at position *i*.

A pooling layer is used to summarize the activations of *P* adjacent neurons by their maximum value

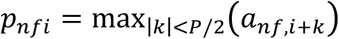

Non-overlapping pooling is applied with step size *p* to decrease the dimension of the input sequence and hence the number of model parameters. The DNA module has multiple pairs of convolutional-pooling layers to learn higher-level interactions between sequence motifs, which are followed by one final fully connected layer with a ReLU activation function. The number of convolutional-pooling layers was optimized on the validation set. For example, two layers were selected for models trained on serum, HCC, and mESC cells, and three layers for the 2i and HepG2 cells (**Additional Table 2**).

### CpG module

The CpG module consists of a non-linear embedding layer to model dependencies between CpG sites *within* cells, which is followed by a bidirectional gated recurrent network (GRU) [24] to model dependencies *between* cells. Inputs are 100*d* vectors *x*_1_,…,*x_T_* where *x_t_* represents the methylation state and distance of *K* = 25 CpG sites to the left and to the right of a target CpG site in cell *t*. Distances were transformed to relative ranges [0; 1] by dividing by the maximum genome-wide distance. The embedding layer is fully connected and transforms *x_t_* into a 256*d* vector 
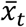
 which allows to learn possible interactions between methylations states and distances within cell *t*:

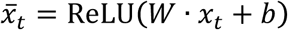

The sequence of vectors 
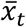
 are then fed into a bidirectional GRU [24], which is a variant of a recurrent neural network (RNN). RNNs have been successfully used for modelling long-range dependencies in natural next [59, 60], acoustic signals [61], and more recently genomic sequences [62, 63]. A GRU scans input sequence vectors 
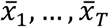
 from left to right, and encodes them into fixed-sized hidden state vectors *h*_1_,…,*h_T_*:

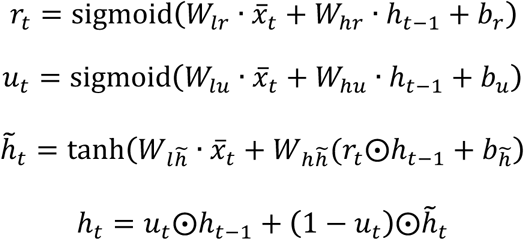

The reset gate *r_t_* and update gate *u_t_* determine the relative weight of the previous hidden state *h_t−1_*and the current input 
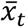
 for updating the current hidden state *h_t_*. The last hidden state *h_T_* summarizes the entire sequence as a fixed-sized vector. Importantly, the set of parameters *W*and *b* are independent of the sequence length *T*, which allows summarising the methylation neighbourhood of a variable number of cells.

To encode cell-to-cell dependencies independently of the order of cells, the CpG module is based on a bidirectional GRU. It consists of a forward and backward GRU with 256*d* hidden state vectors *h_t_*, which scan the input sequence from the left and right, respectively. The last hidden state vector of the forward and backward GRU are concatenated into a 512*d* vector, which froms the output of the CpG module.

### Fusion module

The fusion model takes as input the concatenated last hidden vectors of the DNA and CpG module, and models interactions between the extracted DNA sequence and CpG neighbourhood features via two fully connected hidden layers with 512 neurons and ReLU activation function. The output layer contains *T* sigmoid neurons to predict the methylation rate 
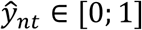
of CpG site *n* in cell *t*:

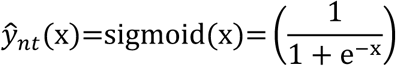

### Model training

Model parameters were learnt on the training set by minimizing the following loss function:

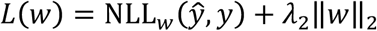

Here, the weight-decay hyper-parameter λ_2_ penalises large model weights quantified by the L2 norm, and 
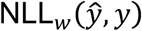
 denotes the negative log-likelihood, which measures how well the predicted methylation rates 
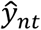
 fit to the true binary methylation states *y_nt_* ∈ {0,1}:

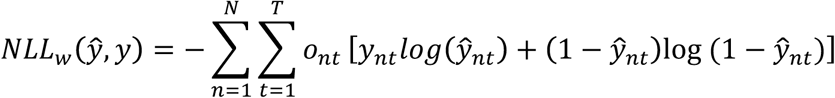

*O_nt_* is one if the true methylation state *y_nt_* is observed for CpG site *n* in cell *t*, and zero otherwise. Dropout [64] with different dropout rates for the sequence, CpG, and fusion module was used for additional regularization. Model parameters were initialized randomly following the approach in Golorot et al. [65]. The loss function was optimized by mini-batch stochastic gradient descent with a batch size of 128 and a global learning rate of 0.0001. The learning rate was adapted by Adam [66], and decayed by a factor of 0.95 after each epoch. Learning was terminated if the validation loss did not improve over ten consecutive epochs (early stopping). The DNA and CpG module were pre-trained independently to predict methylation from the DNA sequence (DeepCpG Seq) or the CpG neighbourhood (DeepCpG CpG). For training the joint module, only the parameters of the hidden layers and the output layers were optimized, whiling keeping the parameters of the pre-trained DNA and CpG module fixed. Training DeepCpG on 18 serum mESCs using a single NVIDIA Tesla K20 GPU took approximately 24 hours for the DNA module, 12hours for the CpG module, and 4 hours for the fusion module. Model hyper-parameters were optimized on the validation set by random sampling [67], and are summarized in **Additional Table 2**. DeepCpG was implemented in Python using Theano [68] 0.8.2 and Keras [69] 1.1.2

## Prediction performance evaluation

### Data pre-processing

We evaluated DeepCpG on different cell types profiled with scBS-seq [5] and scRRBS-seq [8].

ScBS-seq profiled cells contained 18 serum and 12 2i mouse embryonic stem (mESCs), which were pre-processed as described in Smallwood et al. [5], with reads mapped to the GRCm38 mouse genome. We excluded serum cell RSC27_4 and RSC27_7 since their methylation pattern deviated strongly from the remaining serum cells.

ScRRBS-seq profiled cells were downloaded from GEO (GSE65364) and contained 25 human hepatocellular carcinoma cells (HCCs), 6 human heptoplastoma-derived cells (HepG2), and 6 mESCs. Following Hou et al. [8], HCC cell Ca26 was excluded, as well as CpG sites with less than four read counts. HCC and HepG2 cells were mapped to GRCh38, and mESC cells to GRCm38 using the liftOver tool from the UCSC Genome Browser.

Binary CpG methylation states for both scBS-seq and scRRBS-seq profiled cells were obtained for CpG sites with mapped reads, by defining sites with more methylated than un-methylated read counts as methylated, and un-methylated otherwise.

### Holdout validation

For all prediction experiments and evaluations, we used chromosomes 1, 3, 5, 7, 9, and 11 as training set, chromosomes 2, 4, 6, 8, 10, and 12 as test set, and the remaining chromosomes as validation set (**Additional Table 5**). For each cell type, models were fit on the training set, hyper-parameters optimized on the validation set, and all reported accuracies and interpretations exclusively evaluated on the test set. For computing binary evaluation metrics such as accuracy, F1 score, or MCC score, predicted methylation probabilities greater than 0.5 were rounded to one and zero otherwise.

The prediction performance of DeepCpG was compared with averaging CpG sites either in windows within the same cell (WinAvg), or across cells (CpGAvg). We further evaluated random forest classifiers trained on each cell separately using either features similar to DeepCpG (RF), or genome annotation marks as described in Zhang et al. [12] (RF Zhang).

### Window averaging (WinAvg)

For window averaging, the methylation rate 
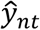
 of CpG site *n* and cell *t* was estimated as the mean of all observed CpG neighbours 
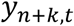
 in a window of length *W = 3,001 bp* centred on the target CpG site *n*:

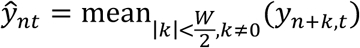

*ŷ_nt_* was set to the mean genome wide methylation rate of cell *t* if no CpG neighbours were present in the window.

### CpG averaging (CpGAvg)

For CpG averaging, the methylation rate 
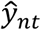
 of CpG site *n* in cell *t* was estimated by averaging the observed methylation states 
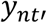 of all other cells 
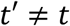
:

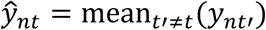

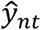
 was set to the genome wide average methylation rate of cell *t* if no methylation states were observed in any of the other cells.

### Random forest models (RF, RF Zhang)

Features of the *RF* model were i) the methylation state and distance of 25 CpG sites to the left and right of the target site (100 features), and ii) *k*-mer frequencies in the *1001 bp genomic* sequence centred on the target site (256 features). The optimal value for K (*k* = 4) was found via hold-out validation (**Additional File 1: Figure 20a**).

The set of features for the *RF Zhang* model (**Additional Table 4**) included a) the methylation state and distance of 2 CpG neighbours to the left and right of the target site (8 features), b) annotated genomic contexts (23 features), c) transcription factor binding sites (24 features), d) histone modification marks (28 features), and e) DNaseI hypersensitivity sites (1 feature). All features were downloaded from the ChipBase database and UCSC Genome Browser for the GRCm37 mouse genome, and mapped to the GRCm38 mouse genome using the liftOver tool from the UCSC Genome Browser.

A separate random forest model was trained for each cell without using data from other cells (**Additional File 1: Figure 20b**). All hyper-parameters, including the number of trees and the tree depth, were optimized for each cell separately on the validation set by random sampling. The implementation is based on the RandomForestClassifer class of the scikit-learn v0.17 Python package.

## Motif analysis

The motif analysis as presented in the main text was performed using the DNA module trained on serum mESCs. Motifs discovered for 2i, HCC, HepG2, and mESC cells are provided in **Additional File 2**. In the following, the filters of the first convolutional layer of the DNA module will be denoted by the motif that they recognize in the input sequence.

### Visualization, motif comparison, GO analysis

Filters of the convolutional layer of the DNA module were visualized by aligning sequence fragments that maximally activated them. Specifically, the activations of all filters were computed for a set of sequences. For each sequence *s_n_* and filter *f* of length *L*, sequence window *s*_*n*,*i*–*L*_, …, *s*_*n*,*i*+*L*_ were selected, if the activation *a_nfi_* of filter *f* at position *i* (Equation 1), was greater than 0.5 of the maximum activation of *f* over all sequences, i.e. *a_nfi_* > 0.5 max*_n,i_*(*a_nfi_*). Selected sequence windows were aligned and visualized as sequence motifs using WebLogo [70] version 3.4.

Motifs discovered by DeepCpG were matched against annotated motifs in the Mus Musculus CIS-BP [43] and UniPROBE [44] database (version 12.12, updated 14 Mar 2016) using Tomtom 4.11.1 from the MEME-Suite [71]. Matches at FDR< 0.05 were reported as significant.

For Genome Ontology (GO) enrichment analysis, the web interface of the GOMo tool of MEME-Suite was used.

### Quantification of motif importance

The importance of each filter was quantified by its activity (occurrence frequency) and its influence on model predictions.

Specifically, the activity of filter *f* for a set of sequences, e.g. within a certain genomic context, was computed as the average of mean sequence activities 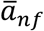
 where 
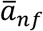
 is the weighted mean of activities *a_nfi_* across all window positions *i* (**
Equation 1
**). A linear weighting function was used to compute 
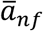
 that assigns the highest relative weight to the centre position.

The influence of filter *f* on the predicted methylation states 
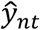
 of cell *t* was computed as the Pearson correlation 
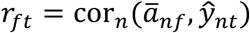
over CpG sites *i*, and the mean influence *r_f_* over all cells by averaging *r_ft_*.

### Motif co-occurrence

The co-occurrence of filters (Figure 3a, **Additional File 1: Additional Figure 9**) was quantified using principal component analysis on the mean sequence activations 
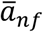

### Conservation analysis

The association between filter activities 
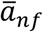
 and sequence conservation was assessed by the Pearson correlation. *PhastCons* [72] conservation scores forthe Glire subset (phastCons60wayGlire) were downloaded from the UCSC Web Browser were used to quantify sequence conservation.

## Effect of sequence and methylation state changes

We used gradient-based optimization as described in Simonyan et al. [56] to quantify the effect of changes in the input sequence *s_n_* on predicted methylation rates 
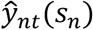
 Specifically, let 
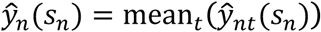
 be the mean predicted methylation rate across cells *t*. Then the effect 
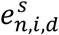
 of changing nucleotide *d* at position *i* was quantified as:

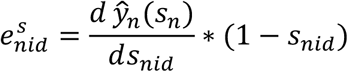

Here, the first term is the first-order gradient of 
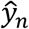
 with respect to *s_nid_*, and the second term sets the effect of wild-type nucleotides ( *s_nid_* = 1) to zero. The overall effect score 
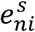
 at position *i* was computed as the maximum absolute effect over all nucleotide changes, i.e. 
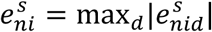
The overall effect of changes at position *i* as shown in Figure 3b was computed as the mean effect 
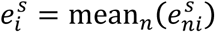
 across all sequences. For the mutation analysis shown in **Additional File 1:Figure 13**, 
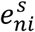
 was correlated with *PhastCons* (phastCons60wayGlire) conservation scores. For quantifying the effect of methylation QTLs (mQTLs) as shown in **Additional 1: Additional Figure 14**, we obtained mQTLs from the supplementary table of Kaplow. et al. [57], and used the DeepCpG DNA module trained on HepG2 cells to compute effect scores. Non-mQTLs variants were randomly sampled within sequence windows, distance-matched to real mQTLs variants.

## Predicting cell-to-cell variability

For predicting cell-to-cell variability (variance) and mean methylation levels, we trained a second neural network with the same architecture as the DNA module, except for the output layer. Specifically, output neurons were replaced by neurons with a sigmoid activation function to predict for a single CpG site *n* both the mean methylation rate 
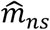
 and cell-to-cell variance 
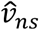
 within a window of size *s* ∈ {1000, 2000, 3000, 4000, 5000} bp. Multiple window sizes were used to make predictions at different scales, using a multi-task architecture, and to mitigate the uncertainty of mean- and variance estimates in low-coverage regions. For training the resulting model, parameters were initialized with the corresponding parameters of the DNA module and fine-tuned, except for motif parameters of the convolutional layer. The training objective was

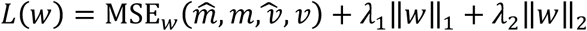

Where MSE the is mean squared error between model predictions and training labels:

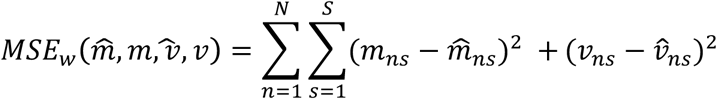

*m_ns_* is the estimated mean methylation level for a window centred on target site *n* of a certain size indexed by *s*:

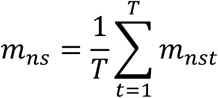

Here,*m_nst_* denotes the estimated mean methylation rate of cell *t* computed by averaging the binary methylation state *y_it_* of all observed CpG sites *Y_nst_* in window *s*:

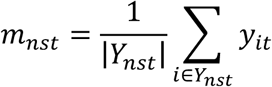

*v_ns_* denotes the estimated cell-to-cell variance

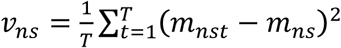

### Identifying motifs associated with cell-to-cell variability

The influence 
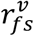
 of filter *f* on cell-to-cell variability in widows of size *s* was computed as the Pearson correlation between mean sequence filter activities 
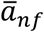
 and predicted variance levels 
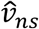
 of sites *n*:

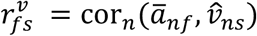

The influence 
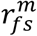 on predicted mean methylation levels 
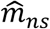
 was computed analogously. The difference 
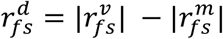
between the absolute value of the influence on variance and mean methylation levels was used to differentiate between motifs that were associated with either high cell-to-cell variance 
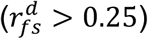
or changes in mean methylation levels 
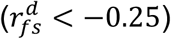

### Functional validation of predicted variability

For functional validation, methylation-transcriptome linkages as reported in Angermueller et al. [10] were correlated with the predicted cell-to-cell variability. Specifically, let 
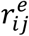
 be the linkage between expression levels of gene *i* and the mean methylation levels of an adjacent region j[10]. Then we correlated 
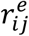
 with *v_j_*, which is the average predicted variability over all CpG sites within context, and FDR adjusted p-values over genes *i* and contexts *j*.

## References

1. Robertson KD. DNA methylation and human disease. Nat. Rev. Genet. 2005;6:597–610.

2. Suzuki MM, Bird A. DNA methylation landscapes: provocative insights from epigenomics. Nat. Rev. Genet. 2008;9:465–76.

3. Laird PW. Principles and challenges of genome-wide DNA methylation analysis. Nat. Rev. Genet. 2010;11:191–203.

4. Jones PA. Functions of DNA methylation: islands, start sites, gene bodies and beyond. Nat. Rev. Genet. 2012;13:484–92.

5. Smallwood SA, Lee HJ, Angermueller C, Krueger F, Saadeh H, Peat J, et al. Single-cell genome-wide bisulfite sequencing for assessing epigenetic heterogeneity. Nat. Methods. 2014;11:817–20.

6. Farlik M, Sheffield NC, Nuzzo A, Datlinger P, Schönegger A, Klughammer J, et al. Single-Cell DNA Methylome Sequencing and Bioinformatic Inference of Epigenomic Cell-State Dynamics. Cell Rep. 2015;10:1386–97.

7. Guo H, Zhu P, Wu X, Li X, Wen L, Tang F. Single-cell methylome landscapes of mouse embryonic stem cells and early embryos analysed using reduced representation bisulfite sequencing. Genome Res. 2013;23:2126–35.

8. Hou Y, Guo H, Cao C, Li X, Hu B, Zhu P, et al. Single-cell triple omics sequencing reveals genetic, epigenetic, and transcriptomic heterogeneity in hepatocellular carcinomas. Cell Res. 2016;26:304–19.

9. Peat JR, Dean W, Clark SJ, Krueger F, Smallwood SA, Ficz G, et al. Genome-wide Bisulfite Sequencing in Zygotes Identifies Demethylation Targets and Maps the Contribution of TET3 Oxidation. Cell Rep. 2014; 9: 1990–2000.

10. Angermueller C, Clark SJ, Lee HJ, Macaulay IC, Teng MJ, Hu TX, et al. Parallel single-cell sequencing links transcriptional and epigenetic heterogeneity. Nat. Methods. 2016; 13: 229–32.

11. Hu Y, Huang K, An Q, Du G, Hu G, Xue J, et al. Simultaneous profiling of transcriptome and DNA methylome from a single cell. Genome Biol. 2016.

12. Zhang W, Spector TD, Deloukas P, Bell JT, Engelhardt BE. Predicting genome-wide DNA methylation using methylation marks, genomic position, and DNA regulatory elements. Genome Biol. 2015; 16: 14.

13. Stevens M, Cheng JB, Li D, Xie M, Hong C, Maire CL, et al. Estimating absolute methylation levels at single-CpG resolution from methylation enrichment and restriction enzyme sequencing methods. Genome Res. 2013; 23: 1541–53.

14. Ernst J, Kellis M. Large-scale imputation of epigenomic datasets for systematic annotation of diverse human tissues. Nat. Biotechnol. 2015; 33: 364–76.

15. Liu Z, Xiao X, Qiu W-R, Chou K-C. iDNA-Methyl: Identifying DNA methylation sites via pseudo trinucleotide composition. Anal. Biochem. 2015; 474: 69–77.

16. Whitaker JW, Chen Z, Wang W. Predicting the human epigenome from DNA motifs. Nat. Methods. 2015; 12: 265–72.

17. LeCun Y, Boser B, Denker JS, Henderson D, Howard RE, Hubbard W, et al. Backpropagation Applied to Handwritten Zip Code Recognition. Neural Comput. 1989; 1: 541–51.

18. Bengio Y. Learning Deep Architectures for AI. 2008;

19. LeCun Y, Bengio Y, Hinton G. Deep learning. Nature. 2015; 521: 436–44.

20. Bhasin M, Zhang H, Reinherz EL, Reche PA. Prediction of methylated CpGs in DNA sequences using a support vector machine. FEBS Lett. 2005; 579: 4302–8.

21. Lu L. Predicting DNA methylation status using word composition. J. Biomed. Sci. Eng. 2010; 03: 672–6.

22. Zhou X, Li Z, Dai Z, Zou X. Prediction of methylation CpGs and their methylation degrees in human DNA sequences. Comput. Biol. Med. 2012; 42: 408–13.

23. Li Z, Chen L, Lai Y, Dai Z, Zou X. The prediction of methylation states in human DNA sequences based on hexanucleotide composition and feature selection. Anal. Methods. 2014; 6: 1897.

24. Chung J, Gulcehre C, Cho K, Bengio Y. Empirical Evaluation of Gated Recurrent Neural Networks on Sequence Modeling. arXiv. 2014.

25. Jarrett K, Kavukcuoglu K, Ranzato M, LeCun Y. What is the best multistage architecture for object recognition? 2009 IEEE 12th Int. Conf. Comput. Vis. 2009. p. 2146–53.

26. Zhang X, Zhao J, LeCun Y. Character-level Convolutional Networks for Text Classification. arXiv. 2015.

27. He K, Zhang X, Ren S, Sun J. Deep Residual Learning for Image Recognition. arXiv. 2015.

28. Szegedy C, Ioffe S, Vanhoucke V. Inception-v4, Inception-ResNet and the Impact of Residual Connections on Learning. arXiv. 2016.

29. Denas O, Taylor J. Deep modeling of gene expression regulation in an erythropoiesis model. Represent. Learn. ICML Workshop. 2013.

30. Alipanahi B, Delong A, Weirauch MT, Frey BJ. Predicting the sequence specificities of DNA- and RNA-binding proteins by deep learning. Nat. Biotechnol. 2015; 33: 831–8.

31. Zhou J, Troyanskaya OG. Predicting effects of noncoding variants with deep learning-based sequence model. Nat Methods. 2015; 12: 931–4.

32. Xiong HY, Alipanahi B, Lee LJ, Bretschneider H, Merico D, Yuen RKC, et al. The human splicing code reveals new insights into the genetic determinants of disease. Science. 2015; 347: 1254806.

33. Kelley DR, Snoek J, Rinn J. Basset: Learning the regulatory code of the accessible genome with deep convolutional neural networks. bioRxiv. 2015.

34. Stormo GD, Schneider TD, Gold L, Ehrenfeucht A. Use of the “ Perceptron” algorithm to distinguish translational initiation sites in E. coli. Nucleic Acids Res. 1982; 10: 2997–3011.

35. Sinha S. On counting position weight matrix matches in a sequence, with application to discriminative motif finding. Bioinformatics. 2006; 22: e454–63.

36. Smallwood SA, Lee HJ, Angermueller C, Krueger F, Saadeh H, Peat J, et al. Single-cell genome-wide bisulfite sequencing for assessing epigenetic heterogeneity. Nat. Methods. 2014; 11: 817–20.

37. Breiman L. Random forests. Mach. Learn. 2001; 45: 5–32.

38. Powers DM. Evaluation: from Precision, Recall and F-measure to ROC, Informedness, Markedness and Correlation. J. Mach. Learn. Technol. 2011; 2: 37–63.

39. Matthews BW. Comparison of the predicted and observed secondary structure of T4 phage lysozyme. Biochim. Biophys. Acta BBA - Protein Struct. 1975; 405: 442–51.

40. Cleveland WS. Robust Locally Weighted Regression and Smoothing Scatterplots. J. Am. Stat. Assoc. 1979; 74: 829–36.

41. Thomson JP, Skene PJ, Selfridge J, Clouaire T, Guy J, Webb S, et al. CpG islands influence chromatin structure via the CpG-binding protein Cfp1. Nature. 2010; 464: 1082–6.

42. Mendenhall EM, Koche RP, Truong T, Zhou VW, Issac B, Chi AS, et al. GC-Rich Sequence Elements Recruit PRC2 in Mammalian ES Cells. Madhani HD, editor. PLoS Genet. 2010; 6: e1001244.

43. Weirauch MT, Yang A, Albu M, Cote AG, Montenegro-Montero A, Drewe P, et al. Determination and Inference of Eukaryotic Transcription Factor Sequence Specificity. Cell. 2014; 158: 1431–43.

44. Newburger DE, Bulyk ML. UniPROBE: an online database of protein binding microarray data on protein-DNA interactions. Nucleic Acids Res. 2009; 37: D77–82.

45. Hervouet E, Vallette FM, Cartron P-F. Dnmt3/transcription factor interactions as crucial players in targeted DNA methylation. Epigenetics. 2009; 4: 487–99.

46. Luu P-L, Scholer HR, Arauzo-Bravo MJ. Disclosing the crosstalk among DNA methylation, transcription factors, and histone marks in human pluripotent cells through discovery of DNA methylation motifs. Genome Res. 2013; 23: 2013–29.

47. Kim TH, Abdullaev ZK, Smith AD, Ching KA, Loukinov DI, Green RD, et al. Analysis of the Vertebrate Insulator Protein CTCF-Binding Sites in the Human Genome. Cell. 2007; 128: 1231–45.

48. Tsai S-Y, Opavsky R, Sharma N, Wu L, Naidu S, Nolan E, et al. Mouse development with a single E2F activator. Nature. 2008; 454: 1137–41.

49. Fernandez-Zapico ME, Lomberk GA, Tsuji S, DeMars CJ, Bardsley MR, Lin Y-H, et al. A functional family-wide screening of SP/KLF proteins identifies a subset of suppressors of KRAS -mediated cell growth. Biochem. J. 2011; 435: 529–37.

50. Lee CS, Sund NJ, Behr R, Herrera PL, Kaestner KH. Foxa2 is required for the differentiation of pancreatic α-cells. Dev. Biol. 2005; 278: 484–95.

51. Wan H, Dingle S, Xu Y, Besnard V, Kaestner KH, Ang S-L, et al. Compensatory Roles of Foxa1 and Foxa2 during Lung Morphogenesis. J. Biol. Chem. 2005; 280: 13809–16.

52. Marais R, Wynne J, Treisman R. The SRF accessory protein Elk-1 contains a growth factor-regulated transcriptional activation domain. Cell. 1993; 73: 381–93.

53. Arsenian S, Weinhold B, Oelgeschläger M, Rüther U, Nordheim A. Serum response factor is essential for mesoderm formation during mouse embryogenesis. EMBO J. 1998; 17: 6289–99.

54. Quenneville S, Verde G, Corsinotti A, Kapopoulou A, Jakobsson J, Offner S, et al. In embryonic stem cells, ZFP57/KAP1 recognize a methylated hexanucleotide to affect chromatin and DNA methylation of imprinting control regions. Mol. Cell. 2011; 44: 361–72.

55. Huang G, Yuan M, Zhang J, Li J, Gong D, Li Y, et al. IL-6 mediates differentiation disorder during spermatogenesis in obesity-associated inflammation by affecting the expression of Zfp637 through the SOCS3/STAT3 pathway. Sci. Rep. 2016; 6: 28012.

56. Simonyan K, Vedaldi A, Zisserman A. Deep Inside Convolutional Networks: Visualising Image Classification Models and Saliency Maps. arXiv. 2013.

57. Kaplow IM, MacIsaac JL, Mah SM, McEwen LM, Kobor MS, Fraser HB. A pooling-based approach to mapping genetic variants associated with DNA methylation. Genome Res. 2015; gr.183749.114.

58. Sumoy L, Carim L, Escarceller M, Nadal M, Gratacòs M, Pujana MA, et al. HMG20A and HMG20B map to human chromosomes 15q24 and 19p13.3 and constitute a distinct class of HMG-box genes with ubiquitous expression. Cytogenet. Genome Res. 2000; 88: 62–7.

59. Bahdanau D, Cho K, Bengio Y. Neural machine translation by jointly learning to align and translate. arXiv. 2014.

60. Wu Y, Schuster M, Chen Z, Le QV, Norouzi M, Macherey W, et al. Google’ s Neural Machine Translation System: Bridging the Gap between Human and Machine Translation. arXiv. 2016.

61. Graves A, Mohamed A-R, Hinton G. Speech recognition with deep recurrent neural networks. 2013 IEEE Int. Conf. Acoust. Speech Signal Process. ICASSPd. p. 6645–9.

62. Lee B, Lee T, Na B, Yoon S. DNA-Level Splice Junction Prediction using Deep Recurrent Neural Networks. arXiv. 2015.

63. Quang D, Xie X. DanQ: a hybrid convolutional and recurrent deep neural network for quantifying the function of DNA sequences. Nucleic Acids Res. 2016; 44: e107–e107.

64. Srivastava N, Hinton G, Krizhevsky A, Sutskever I, Salakhutdinov R. Dropout: A simple way to prevent neural networks from overfitting. J. Mach. Learn. Res. 2014; 15: 1929–58.

65. Glorot X, Bengio Y. Understanding the difficulty of training deep feedforward neural networks. Int. Conf. Artif. Intell. Stat. 2010.

66. Kingma D, Ba J. Adam: A Method for Stochastic Optimization. arXiv. 2014.

67. Bergstra J, Bengio Y. Random search for hyper-parameter optimization. J. Mach. Learn. Res. 2012; 13: 281–305.

68. Bastien F, Lamblin P, Pascanu R, Bergstra J, Goodfellow I, Bergeron A, et al. Theano: new features and speed improvements. arXiv. 2012.

69. Chollet F. Keras: Theano-based deep learning library [Internet]. Available from:https://github.com/fchollet/keras

70. Crooks GE. WebLogo: A Sequence Logo Generator. Genome Res. 2004; 14: 1188–90.

71. Bailey TL, Boden M, Buske FA, Frith M, Grant CE, Clementi L, et al. MEME Suite: tools for motif discovery and searching. Nucleic Acids Res. 2009; 37: W202–8.

72. Siepel A. Evolutionarily conserved elements in vertebrate, insect, worm, and yeast genomes. Genome Res. 2005; 15: 1034–50.

